# Histone code and higher-order chromatin folding: A hypothesis

**DOI:** 10.1101/085860

**Authors:** Kirti Prakash, David Fournier

## Abstract

Histone modifications alone or in combination are thought to modulate chromatin structure and function; a concept termed histone code. By combining evidence from several studies, we investigated if the histone code can play a role in higher-order folding of chromatin. Firstly using genomic data, we analyzed associations between histone modifications at the nucleosome level. We could dissect the composition of individual nucleosomes into five predicted clusters of histone modifications. Secondly, by assembling the raw reads of histone modifications at various length scales, we noticed that the histone mark relationships that exist at nucleosome level tend to be maintained at the higher orders of chromatin folding. Recently, a high-resolution imaging study showed that histone marks belonging to three of the five predicted clusters show structurally distinct and anti-correlated chromatin domains at the level of chromosomes. This made us think that the histone code can have a significant impact in the overall compaction of DNA: at the level of nucleosomes, at the level of genes, and finally at the level of chromosomes. As a result, in this article, we put forward a theory where the histone code drives not only the functionality but also the higher-order folding and compaction of chromatin.

## Introduction

Chromatin is a complex polymer of DNA, histone proteins and other nonstructural proteins [1–4]. Modifications to DNA and histone proteins can affect both the structure and function of chromatin [5, 6]. For instance, posttranslational modifications on the amino acid tail of histone proteins can determine DNA compaction level and in turn, affect the binding of transcription factors to DNA [7–9]. Many of these modifications can occur in a concerted manner to modulate gene expression. A classic example is phosphorylation of histone 3 serine 10 (H3S10p) which has opposing roles during interphase and mitosis [10]. During interphase, phosphorylation at H3S10 is linked to acetylation at H3K9 and H3K14, relaxing chromatin and increasing the accessibility of various transcription factors [11, 12]. In contrast during mitosis, phosphorylation at H3S10 is linked to enhanced methylation at H3K9 to condense chromatin [13, 14]. Overall, this crosstalk can affect signalling pathways to modulate gene expression but also chromatin accessibility. As a result, the combination of histone marks that can potentially regulate chromatin and gene expression is termed as “histone code” [15, 16].

In this article, we hypothesize that higher order chromatin folding can be built up from simple mathematical rules and that the histone code can provide a robust mechanism for this folding. To study the histone code, we defined a framework to correlate 39 histone marks at individual nucleosome positions. Our result shows that histone modifications can be easily clustered into five different groups [17]. One of the clusters is associated with promoters, while others are related to gene bodies or enriched at repeat regions. This result suggests that the histone code is not only present at nucleosome level but also at the level of genes and regulatory regions. We then relate these findings to a previous study of ours where pachytene chromosomes showed distinct compartmentalization of histone modifications at the level of chromosomes [18, 19]. In light of these observation, we propose that the histone code might play a role at various levels of chromatin folding (nucleosome, gene, and chromosome), hinting at a mechanistic folding of DNA in a hierarchical manner.

## Histone code at nucleosome level

To study the associations between histone marks at the nucleosome level, we designed a strategy to assign histone modifications to a given nucleosome. Firstly, we map the ChIP-seq reads for different histone modifications to version hg18 of the human genome using Bowtie [20]. To obtain the best alignment, we set the seed length equal to read length (36 bps), with only two mismatches allowed. Next, we combine the mapped reads of the different histone modifications into one file and predict nucleosome positions using the Nucleosome Positioning from Sequencing (NPS) method [21]. In NPS, each read is extended to 150 nucleotides (nt) and the central 75 nt are taken to get a better estimate of the signal. The peaks which have significantly different distribution of reads on the two strands are removed because the two strands of the DNA should be sequenced evenly. Furthermore, the peaks whose width is less than 80 nt or greater than 250 nt are removed. Next, all the histone modifications are assigned to the predicted nucleosome positions. We count the total number of reads for a particular modification in the nucleosomal peak region and compare it with the expected number of reads for that chromosome using a Poisson distribution. We assign a histone modification to a nucleosome if the p-value for the read count is less than 10^-3^. This computation results in a binary matrix of 0's and 1's. Finally, we compute the correlation coefficients between marks using this binary matrix to describe the associations between the histone modifications. Corrplot and hclust (for hierarchical clustering) functions (R package) are used to plot the correlation matrix and group the histone modifications into distinct clusters.

**Fig. 1.**
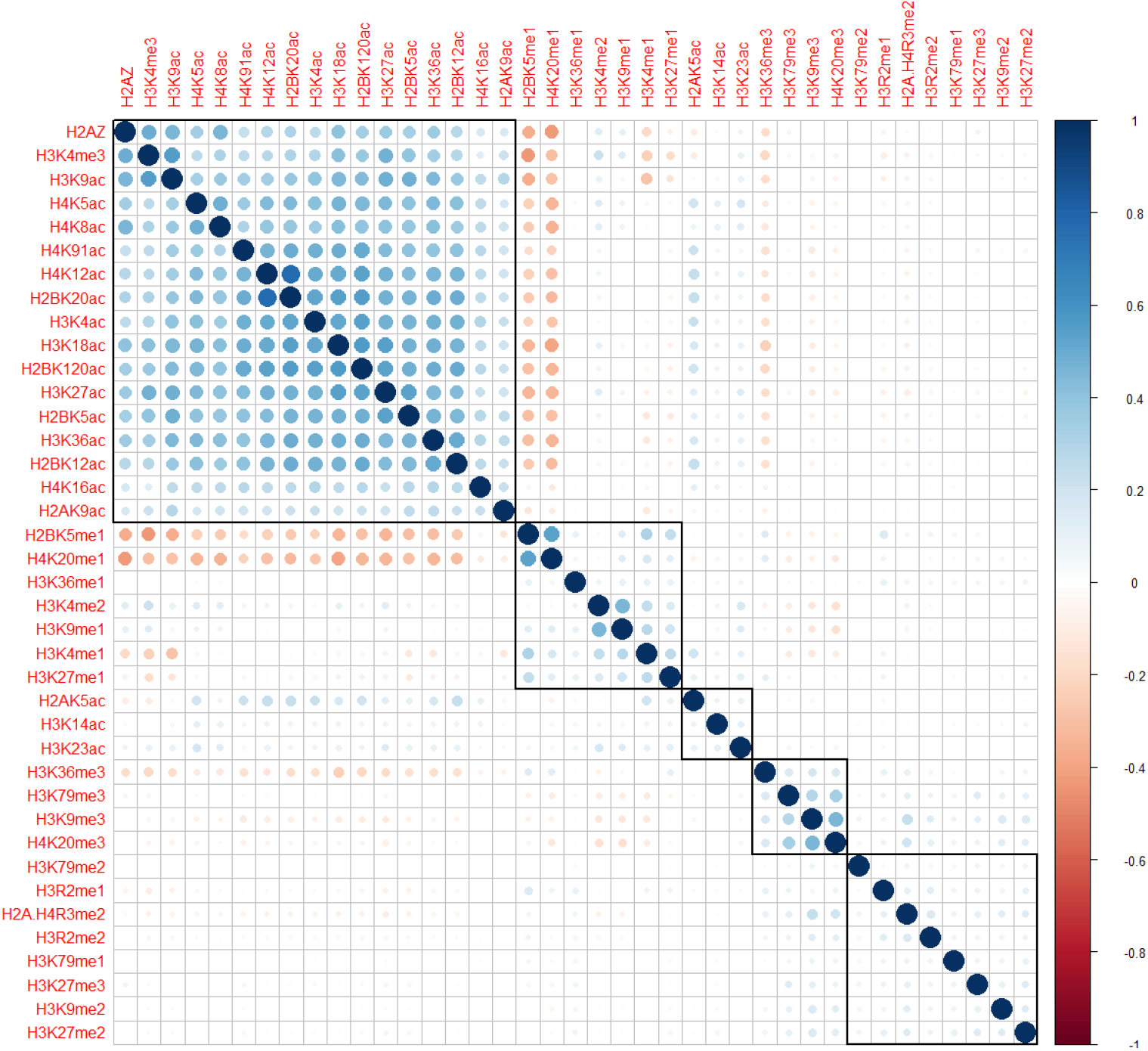
Histone modification associations at the nucleosome level. Five clusters of histone modifications were identified based on 186,492 predicted nucleosome positions using hierarchical clustering. Coefficient of correlation of histone marks at the nucleosome level are presented by a coloring scheme from red (negative correlation) to blue (positive correlation) while white represents an absence of correlation. Black boxes highlight the identified clusters.

## Histone modification associations at nucleosome level are maintained at increasing orders of chromatin folding

We applied our framework to histone modifications data from [22–25], which is still to date the largest collection of histone marks available for a single cell line (CD4 T cells). Our method generated correlation coefficients between histone modifications depicted in Figure 1 using a coloring scheme. The hierarchical clustering identified five clusters of histone modifications. The largest group was found to be associated with active gene expression, comprising acetylation marks and H3K4me3 (Figure 1, top left cluster indicated by a square black box). Another group was associated with inactive chromatin mark (H3K9me3), a mark of constitutive heterochromatin (Figure 1, the fourth cluster from the left). Lastly, we identified a group representative of repressive chromatin (H3K27me3) (Figure 1, the fifth cluster from the left). Using simple magnification of genome browser tracks, we noticed that the histone code which exists at nucleosome level also tends to holds true at the higher order of chromatin folding (Figure 2). In the hypothesis that the histone code does not exist at higher levels of folding, no clustering of histone marks should be visible at higher magnifications, which is not the case here. For instance, the first cluster on the left is observed at promoters and exons while the second group is observed at the gene bodies. This result implies that histone marks associated at the nucleosome level may also pack together at specific sites in the genome at different length scales.

## The histone code defines structural rules at the highest order of chromatin organization

Recently, compartmentalization of active and inactive chromatin marks has been shown in interphase chromosomes [26–29]. However in these studies, a relative spatial positioning of chromatin with respect to a fixed reference is missing. Pachytene chromosomes provide an ideal platform to explore basics of DNA folding, as they are very well defined in 3D space, with a highly reproducible structure. Recently, we published a description of the epigenetic landscape of pachytene chromosomes [18], where we were able to study the relative spatial organization of three histone marks around the central axis of pachytene chromosomes (Figure 3). These histone marks also represent three different clusters in Figure 1. We found these marks to display very different patterns, in terms of size, periodicity and position on the chromosomes (Figure 3). H3K9me3 associated with centromeric chromatin forms helical spreads of length around 500 nm and is located at the end of the chromosome, very close to the central axis (in mouse chromosomes centromeres localize at one of the telomeres [30]). H3K4me3 associated with active chromatin forms 30-60 nm clusters and is located on lateral extensions of the chromosome, probably to facilitate transcription [31–33]. Finally, H3K27me3, a histone mark linked to repressed gene expression [34, 35], shows a remarkable axial periodic and symmetric localization along the chromosomes, hinting for a possible implication in the recombination process [36, 37]. H3K27me3 forms approximately 100-150 nm clusters, which is in between H3K4me3 and H3K9me3 cluster sizes. Overall, this distinct compartmentalization of histone marks hints that histone code might have a structural impact on the overall shape of the chromosomes at the highest order of organization. We summarize our concept of histone code at several levels of hierarchy in a final model (Figure 4).

**Fig. 2.**
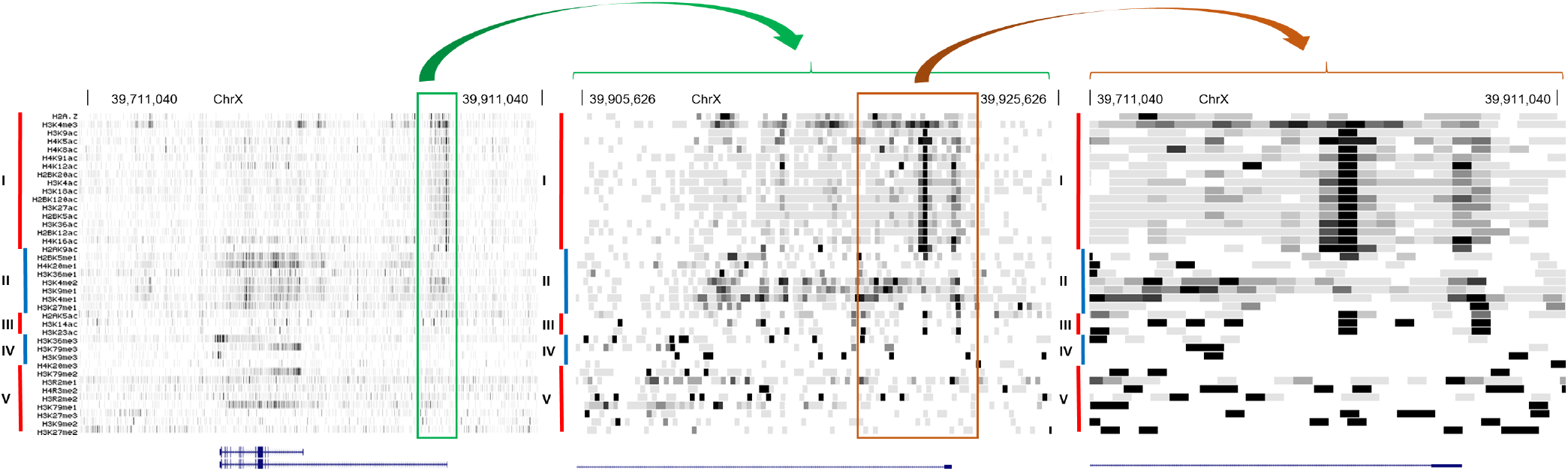
Histone code at lower order tends to be maintained at higher orders. From top to bottom, browser tracks of the different histone modifications shown in the same order as they appear in Figure 1, with clusters displayed on the left (I, II, III, IV and V corresponding to the clusters of Figure 1 in order that appeared top to bottom). Density of reads is represented by a gray scale, with darker regions depicting higher densities. From left to right, three different levels of zoom show the distribution of marks around the locus of gene BCOR. The left panel is the highest magnification at 200 kb, while the right panel is the lowest magnification with a range of 5kb. Genomic positions on X chromosome are indicated on the top of the different graphs for reference. Boxes and arrows in green and orange show the relationship between the three levels of zoom. The reference human genome used is hg18.

## Discussion

Describing the histone code i.e. the functional associations among histone modifications, at different levels of chromatin folding is of particular importance to understand how chromatin structure affects its function. At low order, the histone code can predict if a gene is going to be transcribed or not. At the highest order of chromatin folding, i.e. at the scale of the nucleus, the histone code predicts the general condensation level of chromatin. For instance, H3K9me3 forms large and highly condensed clusters which are located towards the inner side of chromosomes while H3K4me3 forms small decondensed clusters at the periphery of a chromatin domain, most likely to facilitate transcription. Overall, these higher order structures may be transcription units or topologically associated domains (TADs), and may affect the functionality of DNA. For instance, during interphase, regions of active transcription can be brought together in order for transcription to happen [38], while during meiosis regions epigenetically repressed by H3K27me3 are close to the central axis where crossing-overs take place, possibly to silence transcription or prevent repeated regions from being recombined [39].

Nonetheless, histone code is a complex concept and may not hold true all the time [40, 41]. Histone modifications from different clusters can be associated at times, for instance, H3K27me3 and H3K4me3 at poised genes and validating the model will require co-staining of several histone modifications. Furthermore, effort needs to be put to investigate the impact of the histone code on higher-order chromatin folding, using recent genome-wide HiC data [42, 43], and validate these preliminary results.

What we don't know yet is how the three orders, nucleosome, gene and chromosome levels, interact and influence each other. Chromatin looping and folding into particular shapes such as the so-called fractal globule are currently the most advanced explanations on how chromatin can be organised inside the cell nucleus [44–46]. Nonetheless, how a 10-30 nm chromatin fibre transits to commonly observed X-shaped metaphase chromosomes is still an open question.

From our recent studies, we could observe 60 nm clusters of active chromatin domains (H3K4me3), 120 nm of repressed chromatin domains (H3K27me3) and 500 nm of inactive chromatin domains (H3K9me3). From other studies, we know about the existence of 10 nm nucleosomes and 30-100 nm chromatin fibers [47–49]. These 10, 30, 60, 120, 250 nm chromatin domain patterns are highly reminiscent of the 2^*n*^ power law (2, 4, 8, 16, 32, 64, 128, 256, 512, 1024). Using these different measures of clusters as a hint, we hypothesize that chromatin folding could follow a power law of two (at this point, we do not have significant data to support this hypothesis), which is slightly different from the traditional fractal globule model [46]. We propose that the active/inactive state of chromatin is determined by a hierarchical folding design (Figure 4). For instance, the active state of chromatin may only be folded up to the 6th power of 2 i.e. 64 nm but not beyond; increasing levels of compaction will lead to transcriptional repression. This status will be maintained by the modifications from cluster 1 of our histone code table (Figure 1, top left cluster). As a consequence, each of the different levels of chromatin folding, the 10 nm nucleosome, the 30 nm chromatin fiber and the 60 nm nucleosomal domains of H3K4me3 will belong to the active chromatin state. The multiple acetylations and methylations will provide proofreading mechanisms to ensure that this status is maintained. Differently, repressed chromatin will require an additional level of folding, requiring the recruitment of methyltransferases to methylate several sites such as lysine 9 and lysine 27 on histone 3. In this view, a combination of histone modifications is necessary to make sure that no gene is repressed accidentally. For additional levels of folding, a coordinated trimethylation at more specific lysine residues such as lysine 9, lysine 79 on histone 3 and lysine 20 on histone 4 will be required to ensure that the inactive state of chromatin is maintained even after differentiation [50]. Having one active mark to induce or inhibit the transcription of a gene will not be robust. A synergy of modifications or the histone code working in a concerted manner will provide the proper proofreading mechanism to ensure a proper on/off switching of a gene or hierarchical folding of chromatin.

**Fig. 3.**
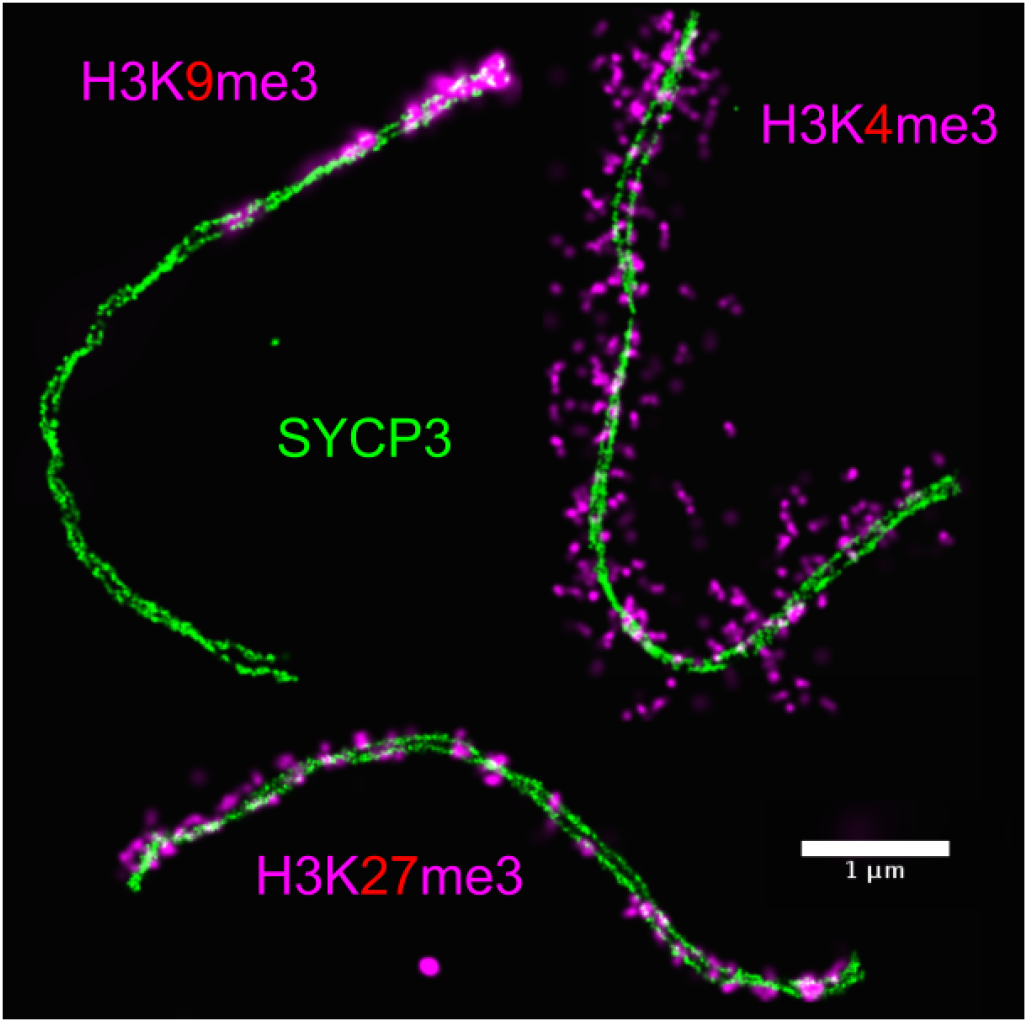
Histone code at the level of chromosomes. Chromatin during pachytene stage of meiosis prophase one is shown to be constrained by anti-correlating distinct clusters of histone modifications (in pink) along the synaptonemal complex (SC) (in green). Active chromatin (H3K4me3) emanates radially in loop-like structures (top right) while repressive chromatin (H3K27me3) is confined to axial regions (bottom) of the SC. Finally, centromeric chromatin (H3K9me3) is found at one end of the SC (top left) [18].

**Fig. 4.**
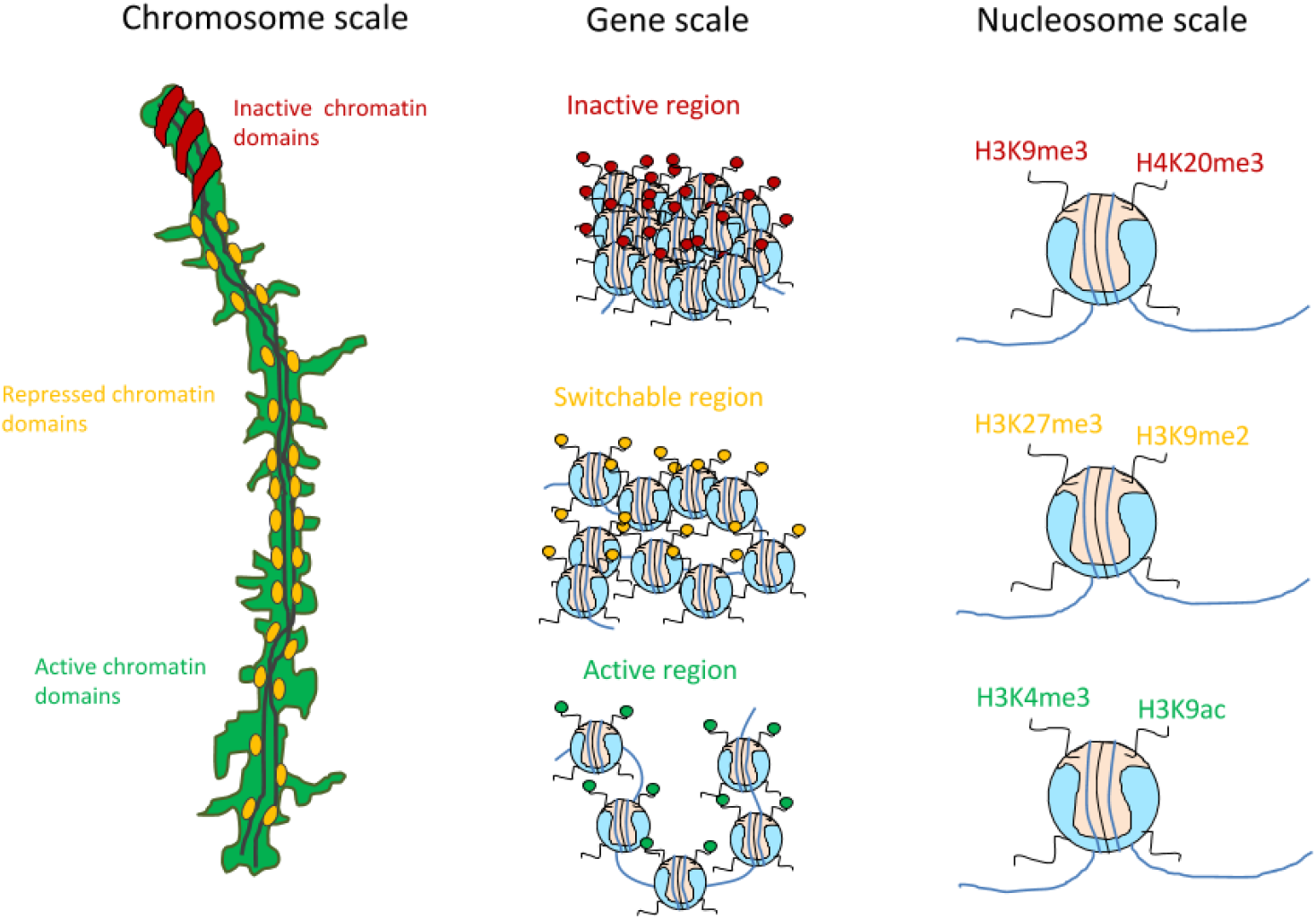
A model for the histone code and its consequences for chromatin structure and function. We propose that the same kind of information (for instance an active mark of chromatin) is clustering at all levels of chromatin folding, in a hierarchical fashion. From left to right, functionally different histone modifications separate in distinct chromatin domains at the chromosome level (left panel, the mouse pachytene chromosomes as a platform to test this hypothesis); separation holds true at the level of genomic regions (middle panel, for instance, an active mark tends to come in arrays/domains); finally information at nucleosome level: histone modifications combinatorially combine to define status of a gene and the future condensation states of chromatin (the so-called histone code, right panel).

## Acknowledgements

K.P. acknowledges Joseph G. Gall and D.F. thanks the Center for Computational Sciences in Mainz for the financial support.

## Author contributions

K.P. initiated and designed the study. K.P. and D.F. performed the analysis and wrote the manuscript.

## References

1. Olins AL, Olins DE (1974) Spheroid chromatin units (*v* bodies). Science 183(4122):330–332.

2. Kornberg RD (1974) Chromatin structure: a repeating unit of histones and dna. Science 184(4139):868–871.

3. Berger SL (2007) The complex language of chromatin regulation during transcription. Nature 447(7143):407–412.

4. Groth A, Rocha W, Verreault A, Almouzni G (2007) Chromatin challenges during dna replication and repair. Cell 128(4):721–733.

5. Klose RJ, Zhang Y (2007) Regulation of histone methylation by demethylimination and demethylation. Nature reviews Molecular cell biology 8(4):307–318.

6. Kouzarides T (2007) Chromatin modifications and their function. Cell 128(4):693–705.

7. Grunstein M (1997) Histone acetylation in chromatin structure and transcription. Nature 389(6649):349–352.

8. Bird AP, Wolffe AP (1999) Methylation-induced repression—belts, braces, and chromatin. Cell 99(5):451–454.

9. Creyghton MP et al. (2010) Histone h3k27ac separates active from poised enhancers and predicts developmental state. Proceedings of the National Academy of Sciences 107(50):21931– 21936.

10. Prigent C, Dimitrov S (2003) Phosphorylation of serine 10 in histone h3, what for? Journal of cell science 116(18):3677–3685.

11. Rea S et al. (2000) Regulation of chromatin structure by site-specific histone h3 methyltransferases. Nature 406(6796):593–599.

12. Rando OJ (2012) Combinatorial complexity in chromatin structure and function: revisiting the histone code. Current opinion in genetics & development 22(2):148–155.

13. Castellano-Pozo M et al. (2013) R loops are linked to histone h3 s10 phosphorylation and chromatin condensation. Molecular cell 52(4):583–590.

14. Fazzio TG, Panning B (2010) Condensin complexes regulate mitotic progression and interphase chromatin structure in embryonic stem cells. The Journal of cell biology 188(4):491– 503.

15. Jenuwein T, Allis CD (2001) Translating the histone code. Science 293(5532):1074–1080.

16. Strahl BD, Allis CD (2000) The language of covalent histone modifications. Nature 403(6765):41–45.

17. Prakash K (2012) A binary combinatorial histone code. Aalto University Master thesis.

18. Prakash K et al. (2015) Superresolution imaging reveals structurally distinct periodic patterns of chromatin along pachytene chromosomes. Proceedings of the National Academy of Sciences 112(47):14635–14640.

19. Prakash K (2016) The periodic and dynamic structure of chromatin. Heidelberg University Phd thesis.

20. Langmead B, Trapnell C, Pop M, Salzberg S (2009) Ultrafast and memory-efficient alignment of short dna sequences to the human genome. Genome Biol 10(3):R25.

21. Zhang Y, Shin H, Song J, Lei Y, Liu X (2008) Identifying positioned nucleosomes with epigenetic marks in human from chip-seq. BMC genomics 9(1):537.

22. Barski A et al. (2007) High-resolution profiling of histone methylations in the human genome. Cell 129(4):823–837.

23. Wang Z et al. (2008) Combinatorial patterns of histone acetylations and methylations in the human genome. Nature genetics 40(7):897–903.

24. Wang Z et al. (2009) Genome-wide mapping of hats and hdacs reveals distinct functions in active and inactive genes. Cell 138(5):1019–1031.

25. Schones D et al. (2008) Dynamic regulation of nucleosome positioning in the human genome. Cell 132(5):887–898.

26. Boettiger AN et al. (2016) Super-resolution imaging reveals distinct chromatin folding for different epigenetic states. Nature 529(7586):418–422.

27. Kirmes I et al. (2015) A transient ischemic environment induces reversible compaction of chromatin. Genome biology 16(1):1–19.

28. Szczurek AT et al. (2014) Single molecule localization microscopy of the distribution of chromatin using hoechst and dapi fluorescent probes. Nucleus 5(4):331–340.

29. ZŻurek-Biesiada D et al. (2016) Localization microscopy of dna in situ using vybrant^®^ dye-cycle” violet fluorescent probe: A new approach to study nuclear nanostructure at single molecule resolution. Experimental cell research 343(2):97–106.

30. Bisig CG et al. (2012) Synaptonemal complex components persist at centromeres and are required for homologous centromere pairing in mouse spermatocytes. PLoS Genet 8(6):e1002701–e1002701.

31. Baudat F et al. (2010) Prdm9 is a major determinant of meiotic recombination hotspots in humans and mice. Science 327(5967):836–840.

32. Baudat F, Imai Y, de Massy B (2013) Meiotic recombination in mammals: localization and regulation. Nature Reviews Genetics 14(11):794–806.

33. Heng H et al. (1996) Regulation of meiotic chromatin loop size by chromosomal position. Proceedings of the National Academy of Sciences 93(7):2795–2800.

34. Arthur RK et al. (2014) Evolution of h3k27me3-marked chromatin is linked to gene expression evolution and to patterns of gene duplication and diversification. Genome research 24(7):1115–1124.

35. Rivera C, Gurard-Levin ZA, Almouzni G, Loyola A (2014) Histone lysine methylation and chromatin replication. Biochimica et Biophysica Acta (BBA)-Gene Regulatory Mechanisms 1839(12):1433–1439.

36. Buard J, Barthés P, Grey C, de Massy B (2009) Distinct histone modifications define initiation and repair of meiotic recombination in the mouse. The EMBO journal 28(17):2616–2624.

37. Hansen KH et al. (2008) A model for transmission of the h3k27me3 epigenetic mark. Nature cell biology 10(11):1291–1300.

38. Cremer T et al. (2015) The 4d nucleome: Evidence for a dynamic nuclear landscape based on co-aligned active and inactive nuclear compartments. FEBS Letters.

39. Termolino P, Cremona G, Consiglio MF, Conicella C (2016) Insights into epigenetic landscape of recombination-free regions. Chromosoma 125(2):301–308.

40. Margueron R, Trojer P, Reinberg D (2005) The key to development: interpreting the histone code? Current opinion in genetics & development 15(2):163–176.

41. Goldberg AD, Allis CD, Bernstein E (2007) Epigenetics: a landscape takes shape. Cell 128(4):635–638.

42. Rao SS et al. (2014) A 3d map of the human genome at kilobase resolution reveals principles of chromatin looping. Cell 159(7):1665–1680.

43. Fortin JP, Hansen KD (2015) Reconstructing a/b compartments as revealed by hi-c using long-range correlations in epigenetic data. Genome biology 16(1):1.

44. Cremer T, Küpper K, Dietzel S, Fakan S (2004) Higher order chromatin architecture in the cell nucleus: on the way from structure to function. Biology of the Cell 96(8):555–567.

45. Albiez H et al. (2006) Chromatin domains and the interchromatin compartment form structurally defined and functionally interacting nuclear networks. Chromosome Research 14(7):707–733.

46. Lieberman-Aiden E et al. (2009) Comprehensive mapping of long-range interactions reveals folding principles of the human genome. science 326(5950):289–293.

47. Gall J (1963) Chromosome fibers from an interphase nucleus. Science 139(3550):120–121.

48. Van Holde K, Zlatanova J (1996) What determines the folding of the chromatin fiber? Proceedings of the National Academy of Sciences 93(20):10548–10555.

49. Zlatanova J, Leuba SH, van Holde K (1999) Chromatin structure revisited. Critical Reviews™ in Eukaryotic Gene Expression 9(3-4).

50. Rada-Iglesias A et al. (2010) A unique chromatin signature uncovers early developmental enhancers in humans. Nature 470(7333):279–283.

